# The variability of amino acids sequences in hepatitis B virus

**DOI:** 10.1101/326959

**Authors:** Jianhao Cao, Shuhong Luo, Yuanyan Xiong

## Abstract

Hepatitis B virus (HBV) is an important human pathogen belonging to the *Hepadnaviridae* family, *Orthohepadnavirus* genus. It infects over 240 million people globally. The reverse transcription during its genome replication leads to low fidelity DNA synthesis, which is the source of variability in the viral proteins. To investigate the variability quantitatively, we retrieved amino acid sequences of 5167 records of all available HBV genotypes (A-J) from the Genbank database. The amino acid sequences encoded by the open reading frames (ORF) S/C/P/X in the HBV genome were extracted and subjected to alignment respectively. We analyzed the variability of the lengths and the sequences of proteins as well as the frequencies of amino acids. Our study comprehensively characterized of the variability and conservation of HBV at the level of amino acids, especially for the structural proteins, hepatitis B surface antigens (HBsAg), to find out the potential sites critical for virus assembly and immune recognition. Interestingly, the preS1/S2 domains in HBsAg were variable at some positions of amino acid residues, which provides a potential mechanism of immune-escape for HBV, while the preS2 and S domains were conserved in the lengths of protein sequences. In the S domain, the cysteine residues and the secondary structures of the alpha-helix and beta-sheet were likely critical for the stable folding of the protein structure. The preC domain and C-terminal domain (CTD) of the core protein are highly conserved. And the polymerases HBpol and the HBx were highly variable at the amino acid level.

## Introduction

Hepatitis B virus (HBV) is one of important human pathogens that causes hepatitis. Over 240 million people are estimated to be infected with HBV. HBV belongs to the *Hepadnaviridae* family, *Orthohepadnavirus* genus. HBV has 3 forms of viral particles, 44nm-diameter infectious Dane particles and non-infectious 22nm-diameter spherical or tubular subviral particles (SVPs). Its genome is only about 3.2kb and contains 4 overlapping opening reading frames (ORFs) which encodes 7 structural or nonstructural proteins. Although HBV is a double-stranded DNA virus, it replicates the genome by an intermediate template of pregenome RNA and thus presents a high mutation rate and low fidelity (1).

There are 3 forms of hepatitis B surface antigens (HBsAg) in the virus envelope with some domains exposed to lumen or cytosol(2). They are large, middle and small HBsAg, respectively and share the same C-terminal domain but different N-terminal domains(3, 4). They form the components of both Dane particles as well as SVPs. The HBsAg can elicit strong protection reaction of individuals against HBV or induce immune tolerance through persistent expression(1, 5). Moreover, it’s a good candidate as therapeutic vaccine but is not efficient enough.

It may provide better information about potential antiviral targets to analyze the variability of HBV at the level of amino acids rather than nucleotide. In this study, to investigate the difference of viral proteins sequences/domains, we reported the sequences variability of HBV strains(genotypes A-J). We also further characterize the features of HBsAg, which may help to understand its function.

## Material and methods

### HBV sequences acquisition and alignment

In November 2017, a total of 5167 HBV genome records with confirmed genotypes were retrieved from the Genbank Nucleotide Database of National Center for Biotechnology Information (NCBI). The whole dataset was divided into different categories according to areas, China, Southeast-Asia (SE-Asia, including Indonesia, Malaysia, Myanmar, South Korea, Thailand, Vietnam, Japan and Korea), America (including Argentina, Brazil, Canada, Chile, Colombia, Mexico, USA and Venezuela), Europe (including Belgium, France, Germany, Ireland, Italy, Luxembourg, Netherlands, Poland, Russia, Serbia, Spain, Sweden, Turkey and UK), and the other area (including India, Iran, Saudi Arabia and Syria). Then, the amino acid sequences of 4 HBV ORFs (S, C, P and X) were extracted according to their start and end sites and were then subjected to sequences alignment with Clustral Omega(6).

Subsequent analysis was focused on those functional sequences or domains annotations. The sequences labelled with “nonfunctional” or “truncated” were discarded. The final items for analysis were 3208 sequences of ORF C, 3064 sequences of ORF S, 3701 sequences of ORF P and 4265 sequences for ORF X. The domains for further analysis were the preS1 receptor binding domain (preS1-RBD), the preS1 domain, the preS2 domain, the S domain, the preC domain, the core protein assembly domain (core-AD, usually 149AAs) and the core protein C-terminal domain (core-CTD, also known as the arginine-rich domain, ARD). The positions of the preS1-RBD is referred to the sequence previously reported(7). The preS1/S2/S domains are encoded by ORF S, while the preC, core-AD and core-CTD are encoded by ORF C.

### Feature of HBV sequences and domains

We analyzed the variability of sequences from different aspects. All domains were extracted from aligned sequences. Aligned sequences with only gaps across the domain were discarded in subsequent analysis. Hence, the left items are the total number of sequences for calculation of the percentage of different indices mentioned below. Among all the sequences or domains the most frequent one was defined as the predominant sequence, which represents the conserved sequence. It was defined as unique sequences that those sequences or domains encodes the same protein but different to each other at the level of amino acid. Similarly, the most frequent length of sequences or domains was defined as the predominant one. The ratio of sequences with the predominant length suggests the length conservation of a sequence or domain.

### Statistics of the frequency of amino acids of each site in sequences

After sequences alignment, we computed the ratio of amino acids residues (including gaps) for each position as below.

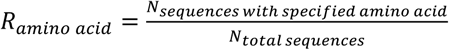

The ratio was the the number of sequences with specified amino acid divided by the total number of available sequences. We define the predominant amino acid residue as the one with the highest ratio at that site.

### Prediction of the secondary structure of HBsAg

A representative sequence record (Accession number of its genome record: AM295797) was subjected to the prediction of secondary structure of the large/middle/small HBsAg. The prediction was performed in PSIPRED website (http://bioinf.cs.ucl.ac.uk/psipred/).

### Analysis of similarities of pair-wise sequences

The pair-wise sequences similarity was calculated based on the aligned sequences. The similarity was defined as the ratio of the number of positions with identical amino acid residues to the length of the sequences. The similarity profile was shown as a histogram. Pearson correlation coefficient (R) was computed between two profiles using Python library scipy. R>0.8 was regarded as correlation, while R>0.9 was regarded as high correlation.

## Results

### Amino acid sequences analysis

The geographical sources of the 5167 sequences of documented HBV strains are listed in Table 1. A total of 3657 items(70.8%) of them are from Asia, 42.2% of which are China strains, and 17.8% are Europe strains. It suggests that China is still a major epidemic area.

**Table 1.** The geographical distribution of 5167 HBV records

| Asia |  | Europe |  | North/South American |  | Africa |  | Oceania |  |
| --- | --- | --- | --- | --- | --- | --- | --- | --- | --- |
| China | 2181 | UK | 349 | USA | 185 | South Africa | 73 | Australia | 24 |
| Japan | 516 | Belgium | 151 | Argentina | 150 | Gabon | 1 | NewZealand | 1 |
| India | 308 | Luxembourg | 124 | Canada | 66 |  |  |  |  |
| Malaysia | 237 | Sweden | 97 | Brazil | 33 |  |  |  |  |
| Vietnam | 134 | Poland | 53 | Chile | 22 |  |  |  |  |
| Iran | 86 | France | 43 | Mexico | 17 |  |  |  |  |
| Syria | 70 | Netherlands | 34 | Venezuela | 13 |  |  |  |  |
| Thailand | 48 | Germany | 22 | Colombia | 4 |  |  |  |  |
| Indonesia | 40 | Italy | 17 |  |  |  |  |  |  |
| Myanmar | 15 | Turkey | 12 |  |  |  |  |  |  |
| South Korea | 10 | Spain | 11 |  |  |  |  |  |  |
| Korea | 6 | Serbia | 5 |  |  |  |  |  |  |
| SaudiArabia | 6 | Ireland | 2 |  |  |  |  |  |  |
|  |  | Russia | 1 |  |  |  |  |  |  |
| **(Total)** | **3657** |  | **921** |  | **490** |  | **74** |  | **25** |

We compared 4 features of HBV sequences, the ratio of the predominant sequence, the ratio of unique sequences, the ratio of sequences with the predominant length and the number of sequence lengths (Fig. 1). These values can characterized either the variability or the conservation of sequences or domains. The preC domain presented the highest ratio of predominant sequence, indicating a highly conserved domain, followed by the core-CTD. The HBpol and large HBsAg had the top two highest ratio of unique sequences.

**Fig. 1.**
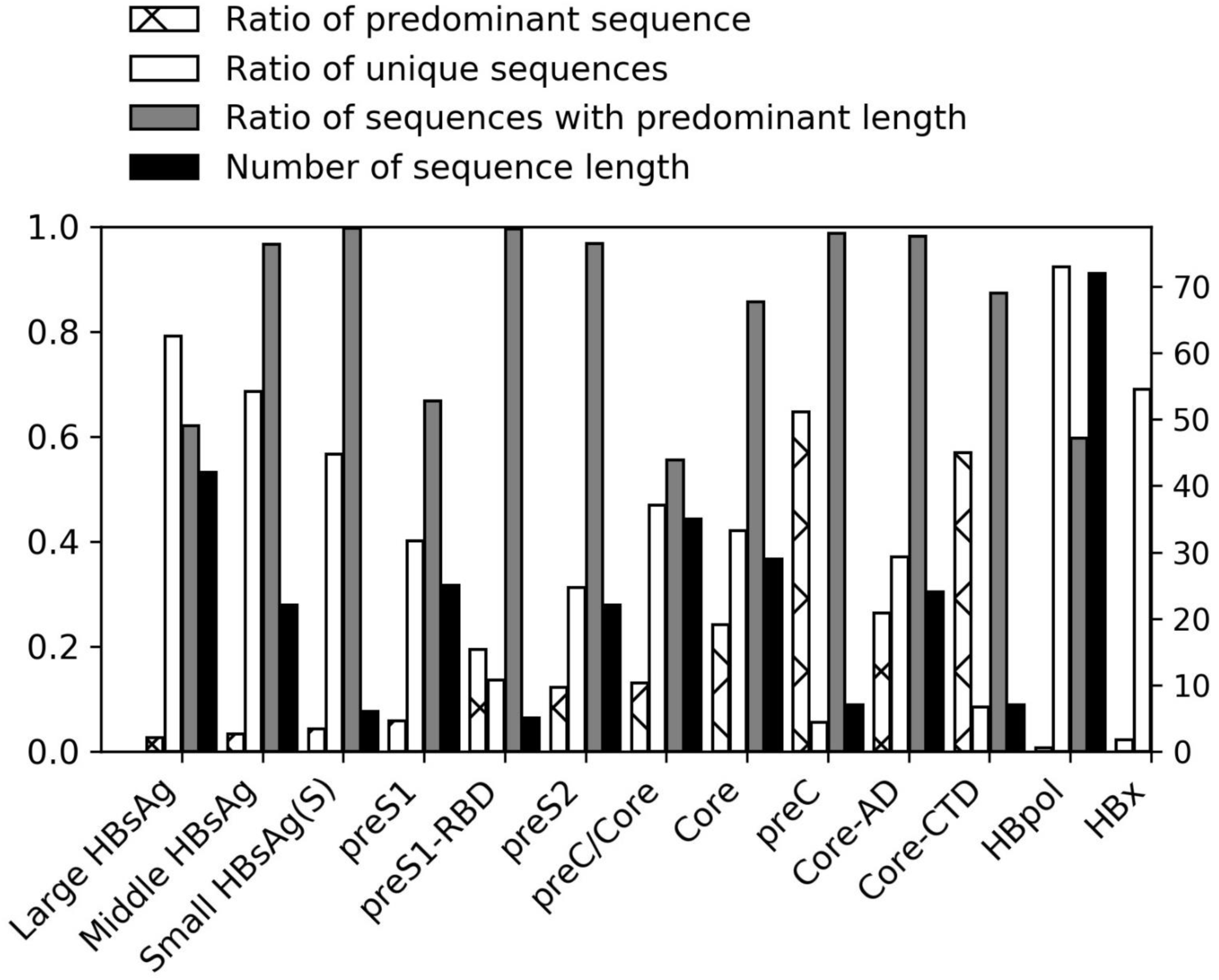
The statistical property of different amino acid sequences/domains in all HBV strains. (a) The ratio of predominant sequence (mesh bar). The ratios of predominant sequence are smaller than 70%, even that of HBcAg is not more than 30%. However, the sequences of the preC domain and the core-CTD are highly conserved. (b) The ratio of all unique sequences (white bar). Higher ratio indicates higher variability of the sequence/domain. (c) The ratio of sequences with predominant length (grey bar). The most adopted length indicates the conserved sequence or domain confines to the length of amino acid residues. The most conserved domains are the preC domain and the core-CTD, over 50% sequences of which have the same sequences. (d) The number of sequence length (black bar). It indicates the variability of sequence/domain in length. It also suggests the small HBsAg, the preC domain and the core-CTD are the most conserved.

The lengths of amino acid sequences or domains of HBV was variable. The number of the lengths of small HBsAg was the smallest (Fig. 1 and Fig. S1), while those of the HBpol was the largest. The preC domain and core-CTD were next to small HBsAg, which revealed their lengths were conserved. The preS1/S2 domains and the HBcAg are the most conserved. However, considering the length, the small and middle HBsAg, the preS1-RBD, the preS2 domain, the core-AD and the HBx are the most conserved. It’s probable that the length is critical for the assembly, so structural proteins, especially the small HBsAg and the core-AD, were conserved except the preS1 region which functions as receptor binding(7). Moreover, for the HBpol, the sequence length aren’t critical. Some domains of the HBpol may provide the flexibility of length for the function.

We analyzed different HBV proteins at the level of amino acid and found that only the predominant sequence of HBcAg account for more than 20% of total sequences (Fig. 1). However, almost all of small HBsAg sequences are of the same length. The preS2 is also highly conserved in length. While the preS1 is more variable. In comparison, the HBpol was variable both in length and in amino acid sequence.

### The predominant AAs in different sequences or domains

We analyzed the amino acid residues in 4 HBV ORFs. As a whole, it’s interesting that some sites were much conserved while some were much variable (Fig. 2). Frequencies of almost all predominant AAs were above 50%. AAs in the ORF S were more variable. For those predominant AAs (>=0.950), there were only about ∼20% sites in the preS1 domain and ∼50% sites in the preS2 domain, while there were more than 60% sites in other sequences or domains, especially near 70% sites in the HBpol. Even in the ORF C, there were more than 60% sites with predominant AAs (>=0.950). The amino acid residues in the preS2 domain and the S domain were more variable than those in the core-AD and the core-CTD. It indicates the surface antigen could be more variable and provides the base of immune escape. Among all sites, L350 in the large HBsAg and H83 in the HBpol were the most conserved. No mutation was found at those sites of all HBV strains in our study. It is probable a potential target for development of new anti-virus strategy.

**Fig. 2.**
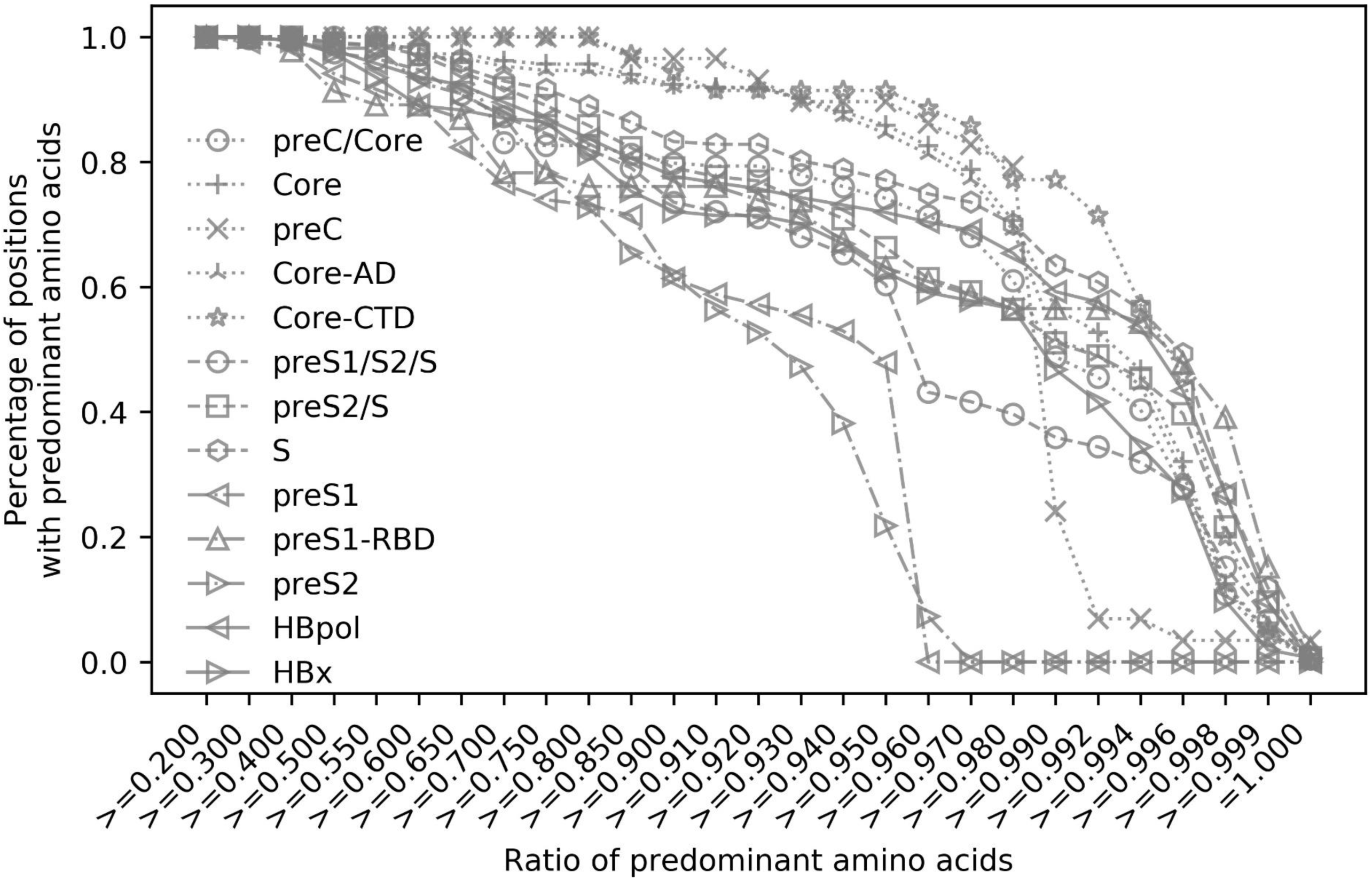
The ratio of sites at different percentage of predominant amino acid. The ratio of sites with high percentage of predominant residues is lower in the preS1/S2 domain than the S domain. It indicates higher variability of the preS1/S2 domains.

### Sequences variant

After sequences alignment, we could further obtain the profile of pair-wise similarity in a whole picture. To compare the similarity of sequences or domains in different areas, we extracted subsets data from different locations, especially China which is a very important area with high prevalence of HBV infection. It’s surprising that different areas exhibit different profiles of pair-wise similarity (Fig. 3). Almost all pair-wise similarities were more than 80% except that some similarities of preS2 were around 60%. The profiles of both the preC domain and the core-CTD in different areas were highly similar. When these profiles were compared using the Pearson cross-correlation coefficient, the preS2 domain, the preC domain and the core-CTD show highly coefficient. However, the profiles of the preS1 domain, the S domain, the core-AD, the HBpol and the HBx were different among different areas.

**Fig. 3.**
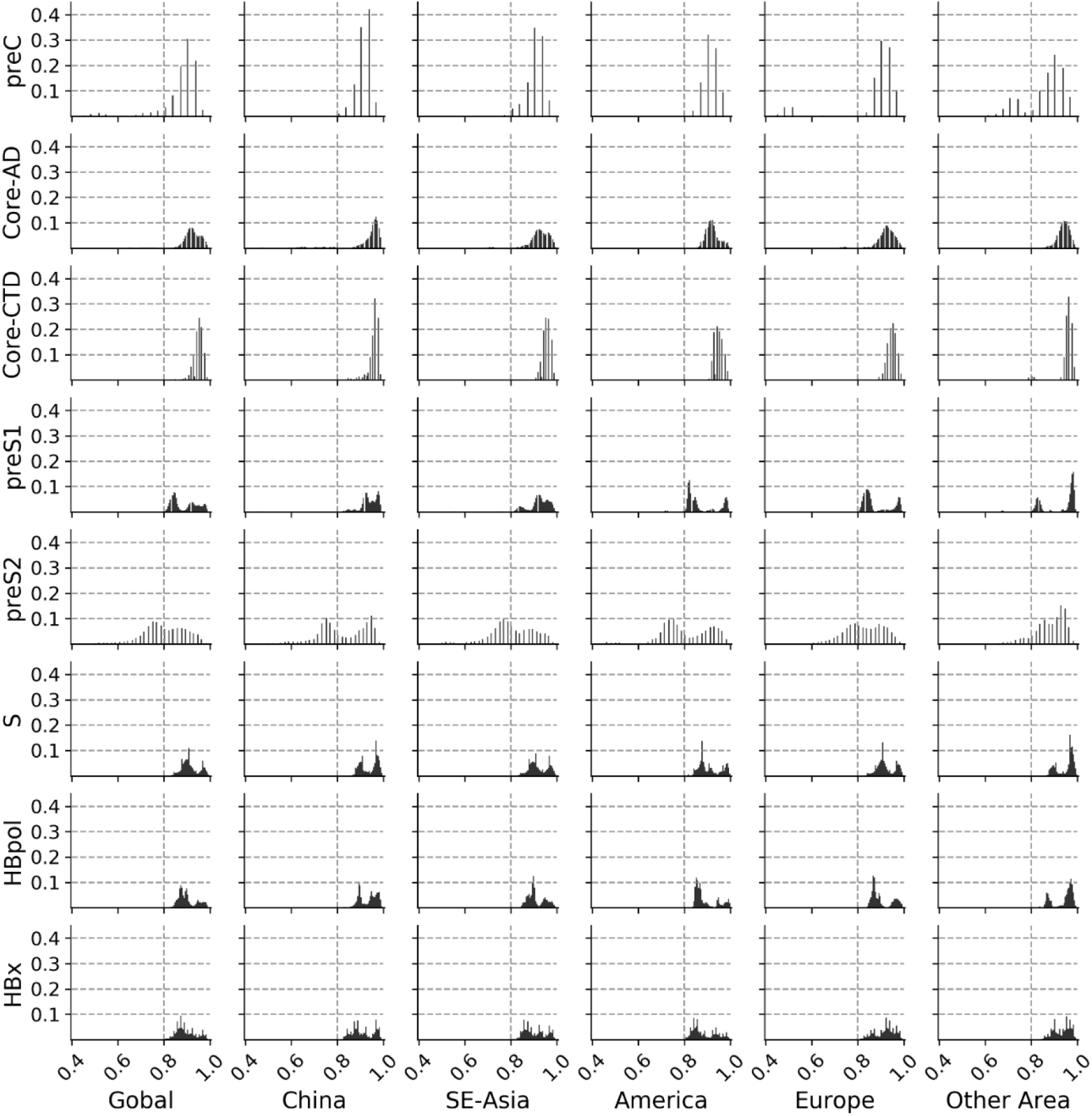
The profile of pair-wise similarity in different sequences/domains of different areas. SE-Asia (Southeast Asia) includes Indonesia, Malaysia, Myanmar, South Korea, Thailand, Vietnam, Japan and Korea. America includes Argentina, Brazil, Canada, Chile, Colombia, Mexico, USA and Venezuela. Europe includes Belgium, France, Germany, Ireland, Italy, Luxembourg, Netherlands, Poland, Russia, Serbia, Spain, Sweden, Turkey and UK. Other areas includes India, Iran, Saudi Arabia and Syria. The sequence number of each area refer to Table 1.

### Prediction of secondary structure of the HBsAg

The HBsAg component are important proteins in immune escape and persistence of immune tolerance. It’s still known a little about the structure of it. From the prediction of its secondary structure, it’s surprising that alpha-helix only exists in the S domain, while the preS1/S2 domain contains only coil (Fig. S3). While comparing the different amino acid frequencies of each sites in the HBsAg, we also found that cysteine residues also exist in the S domain. Considered the conserved length of S domain, we speculate that the stable of its structure is critical for the assembly of both SVPs and envelope of Dane particles. There were 40-46 amino acid residues in the N-terminal of preS1 critical for the receptor binding of HBV(7), so the flexibility in the preS1/S2 domain could provide the allosteric effect during the virus-host recognition.

### The distribution of predominant amino acids in the HBsAg

For the HBsAg sequence, it’s interesting that cysteine only exists in S domain (Fig. S2). While for HBcAg, another structural protein of virions, only 3 highly conserved cysteine residues exist in the core-CTD (Fig. S3). However, cysteine residues exist throughout sequences of the non-structural proteins, the HBpol and the HBx (Fig. S4 and Fig. S5). A special function of cysteine is to form intra- or inter-molecular disulfide bonds, which can stable the structure of proteins. Thus, it’s postulated that these disulfide bonds keep the stability of the S domain in the envelope, while the preS1/S2 domain without any cysteine residues could provide highly flexible conformation to help binding to the host receptor.

It have been reported that 40-46 amino acid residues in the N-terminal of the preS1 domain is critical for binding to host receptor(7-9). We found that this domain is highly conserved. However, it’s surprising that over 50% HBV genome records encodes the preS1 sequence with a N-terminal extension of RBD up to 11 amino acid residues (Fig. 4). It has never been reported that this sequence is associated with receptor binding. Its amino acid sites were also highly conserved excepting Lys11 (Fig. 4).

**Fig. 4.**
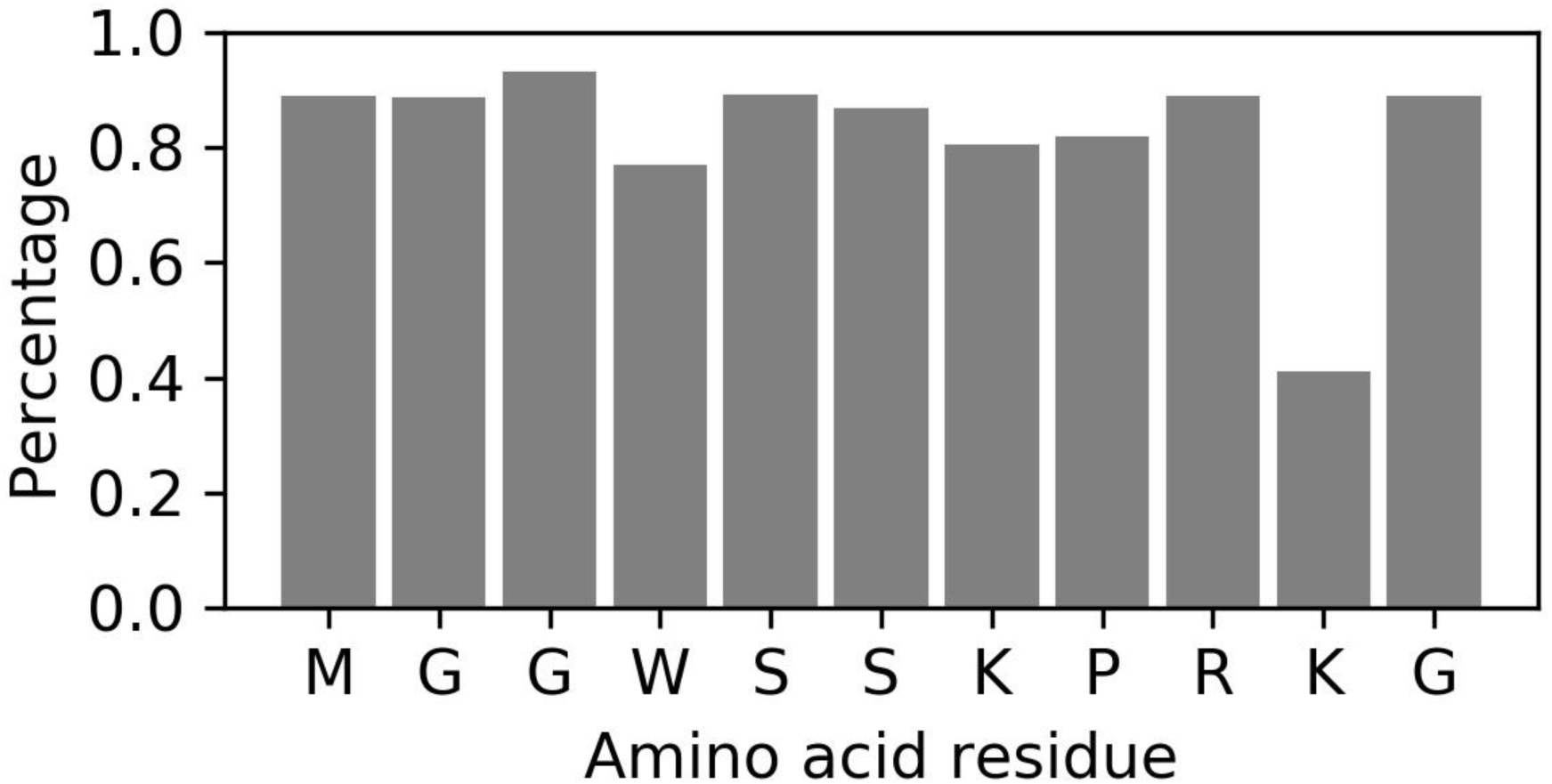
The 11 amino acid residues in the upstream of the preS1-RBD. Most residues were highly conserved except that Lys10.

## Discussion

The amino acid sequence is close to the folding of proteins. It causes high variation that HBV genome replicates through a process of reverse transcription. Thus, it’s necessary to have a comprehensive investigation on the amino acid sequences of HBV proteins, which could help finding the variability and conservation of domains close to the function and the differences between different geographical strains.

Our study revealed the variability of HBV ORF S, C, P and X at the level of amino acid. It showed the sequence variability both in length and in the amino acid sequences. These variability of virus proteins don’t seem to hamper the normal functions. The 4 ORFs of HBV are partially overlapping. When a mutation happens in a ORF, it also likely happens in another one. It will effect either the function or the transcription of viral proteins. Hence, the conserved regions in HBV genome are functionally important. However, it also reveals functional flexibility in high variable regions.

Especially, the variability of the HBsAg at the level of amino acid provided many potential epitopes. This is the potential mechanism exhausting the immunocytes or antibodies which recognize the HBsAg. It has been reported that the HBsAg in tubular SVPs organizes regularly in crystalline-like pattern(10). As the C-terminal part of the HBsAg, the S domain is transmembrane and folds as the protrusion on the periphery(2, 10). Moreover, there were a total of 12 highly conserved cysteine residues in the S domain, far more than those in the core-AD of HBcAg (Fig. S3 and Fig. S4). It indicates that these cysteine residues were probably critical for the stable of protein structure, which could help the protrusions on SVPs to arrange in a regular way. Even the cysteine residues in HBcAg are not necessary for the disulfide bond(11). The core-CTD is located in the internal of HBV capsid, which helps enclosing the virus genome in assembly(12). We postulate that those cysteine residues and the lengths of the S domain are much critical for the stable of the HBsAg structure. Furthermore, the N-terminal extension of the preS1-RBD was also a highly conserved sequence. It’s probably associate with the function of receptor binding in some unknown way.

In conclusion, we studied the viral proteins of HBV at the level of amino acid. Quantitatively investigation reveals the conservation and variability among different sequences and domains. It would be helpful to further study the variant epitopes of the HBsAg in the immune escape and recognition of HBV and for the development of new vaccines and antiviral drugs.

## Acknowledgements

The authors would like to thank Prof. Ping Zhu (Institute of biophysics, Chinese Academy of Sciences) and Prof. Jingqiang Zhang (Sun Yat-sen university) provided help in this research. This work was supported by National Basic Research Program (973 Program 2012CB911201), National Natural Science Foundation of China (NSFC 31171397, 31271533, 31570827)

## Author contributions

JC, SL and YX designed the study. JC conducted computational work, JC and YX performed data analysis. JC, SL and YX wrote the manuscript draft.

## Compliance with ethical standards

**Conflicts of interest.** The authors declare that they have no conflicts of interests.

## Supplement materials

**Fig. S1.**
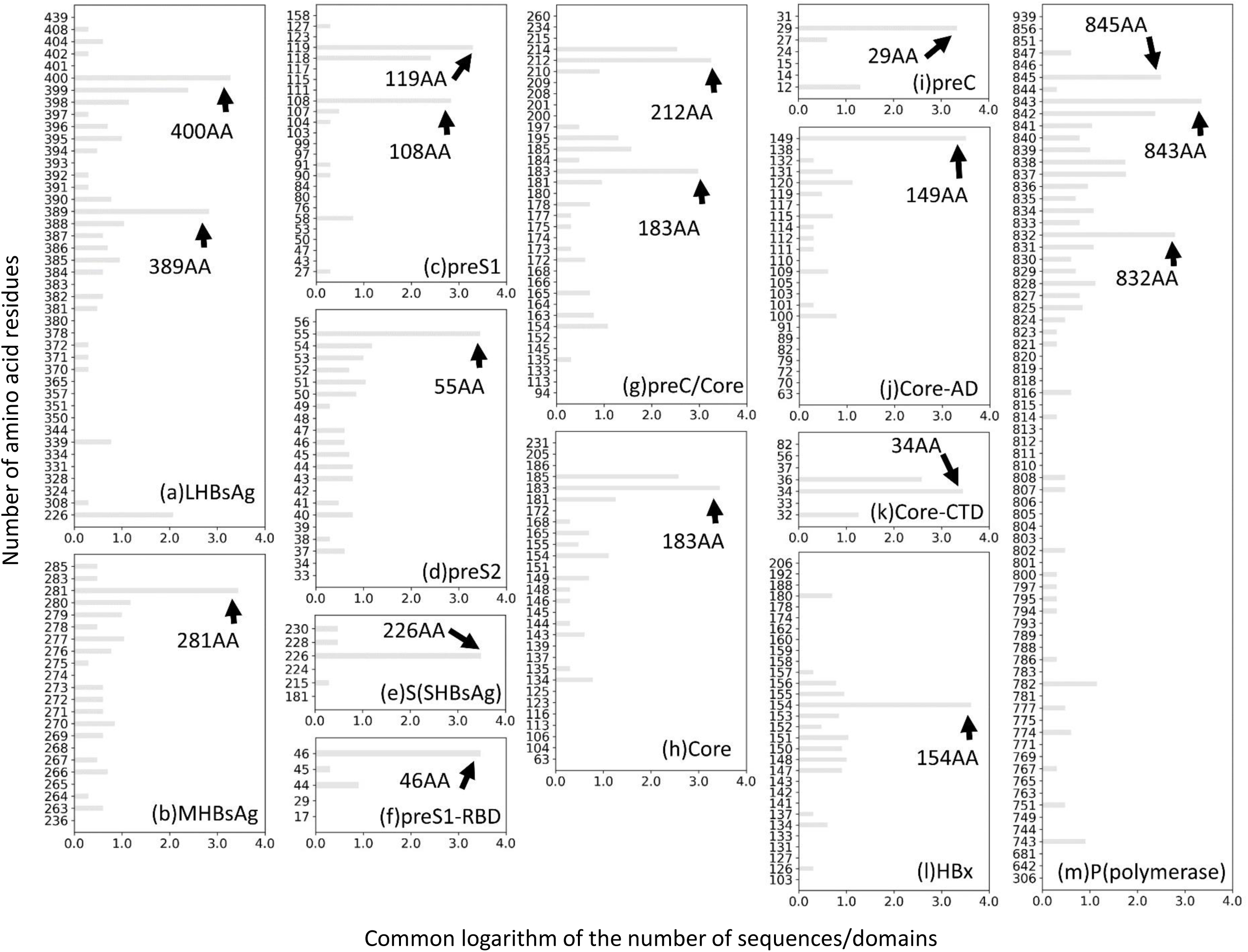
The detail of adopted length by different sequences/domains. The Y-axes is the common logarithm of the number of sequences/domains. The sequences/domains are indicated. (a) the large HBsAg, (b) the middle HBsAg, (c) the preS1 domain, (d) the preS2 domain, (e) the S domain (the small HBsAg), (f)re the preS1-RBD (receptor binding domain), (g) the preC/core, (h) the core domain, (i) the preC domain, (j) the core-AD, (k) the core-CTD, (l) the HBx, (m) P (the polymerase).

**Fig. S2.**
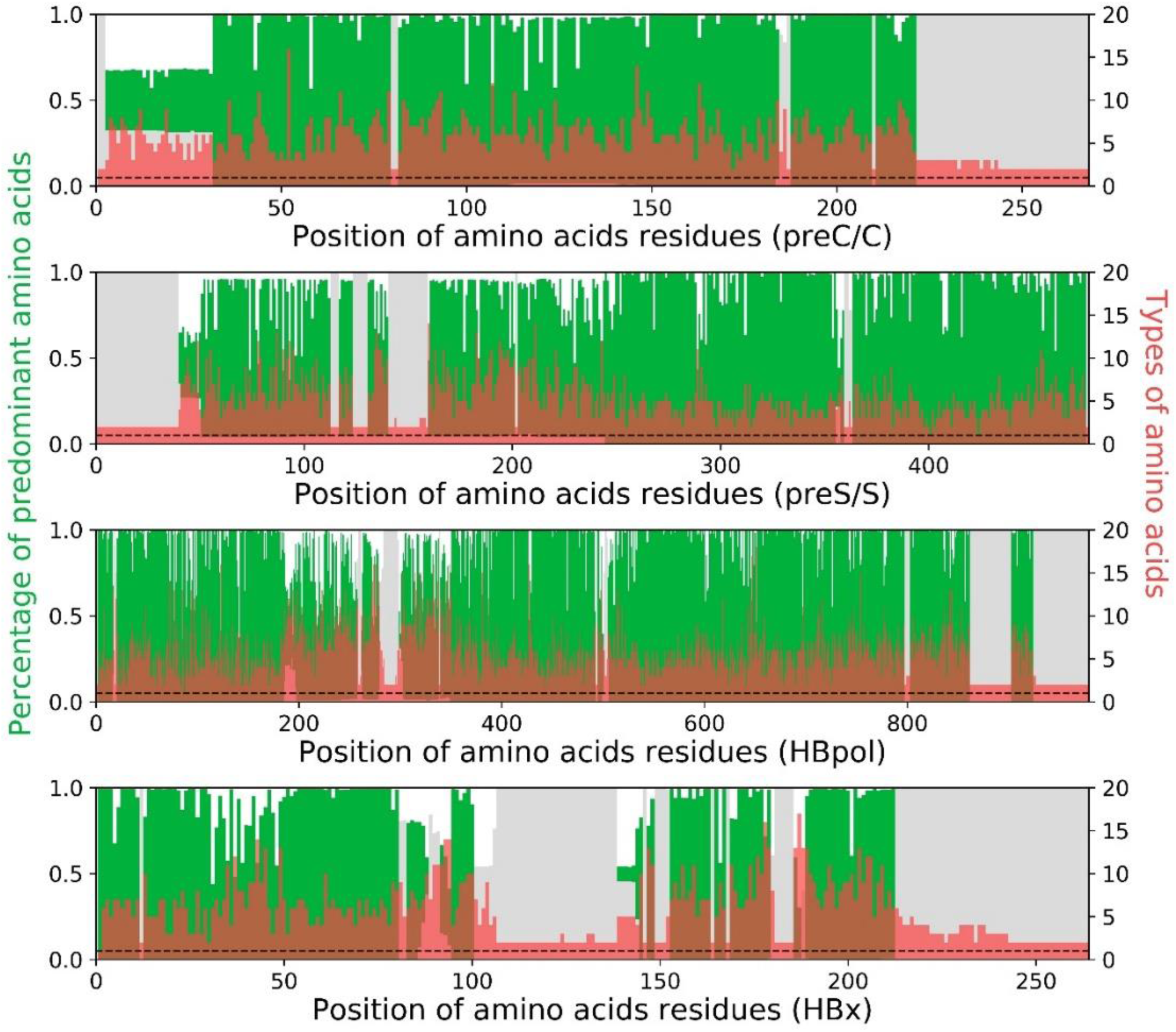
The profile of predominant amino acids in 4 HBV ORFs. Green bar indicates the ratio of sequences of predominant amino acid residue in correspondent sites. While the red bar indicates the types of amino acid residues. Grey bar indicates the ratio of sequences of the gaps after alignment.

**Fig. S3.**
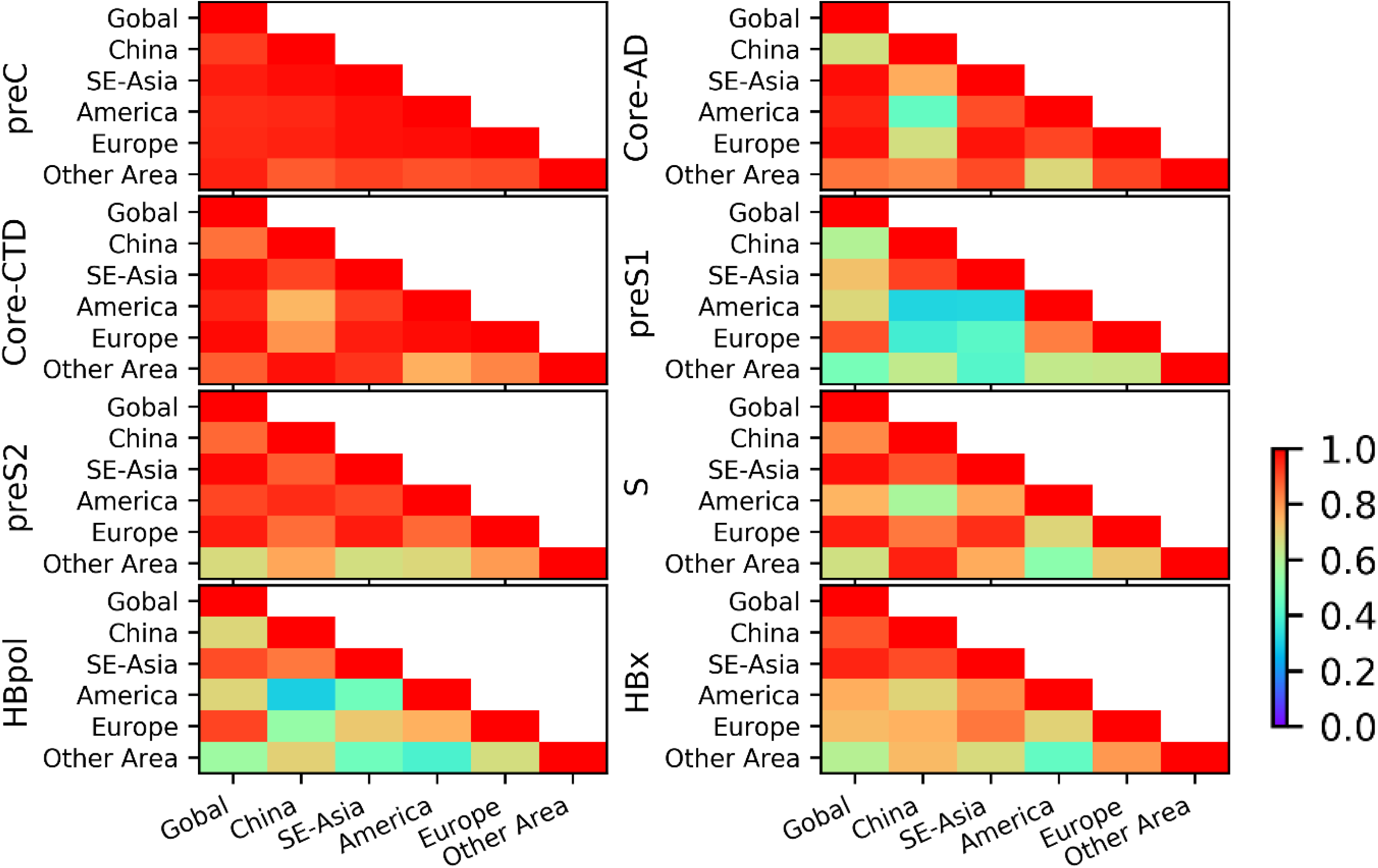
The Pearson cross-correlation coefficient of the profiles of pair-wise similarity in Fig 4 (P<0.001).

**Fig. S4.**
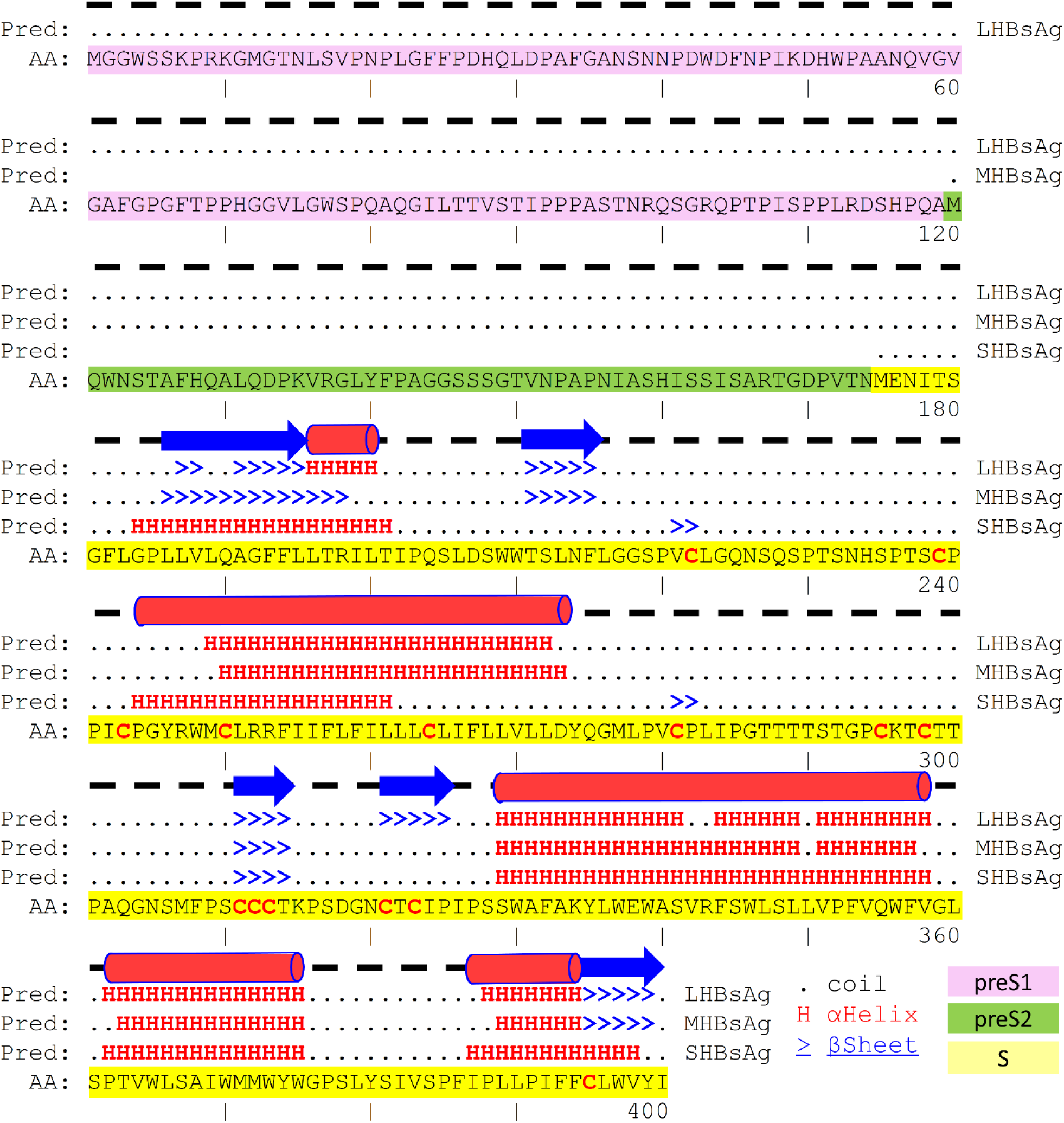
The prediction of the secondary structure. Representation is the prediction of 3 forms of HBsAg components. Alpha-helix and beta-sheet only exist in the S domain. The preS1, preS2 and S domains are colored in purple, green and yellow respectively. Cysteine residues are colored in red.

**Fig. S5.**
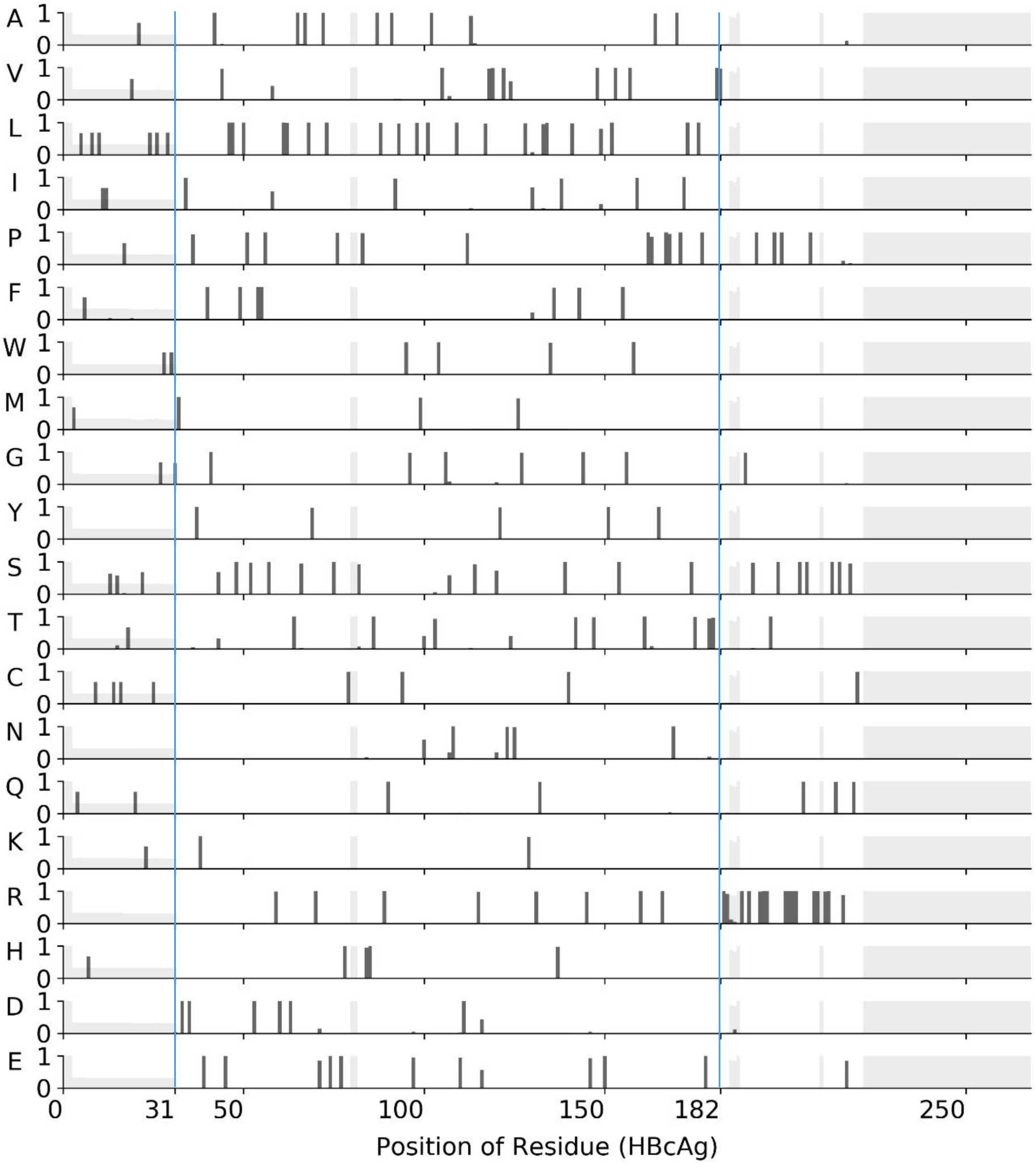
The frequencies of 20 amino acids in different sites of ORF C (preC/HBcAg). The position (1-30) is the preC domain. Position (31-181) is the core-AD, position (182-268) is the core-CTD.

**Fig. S6.**
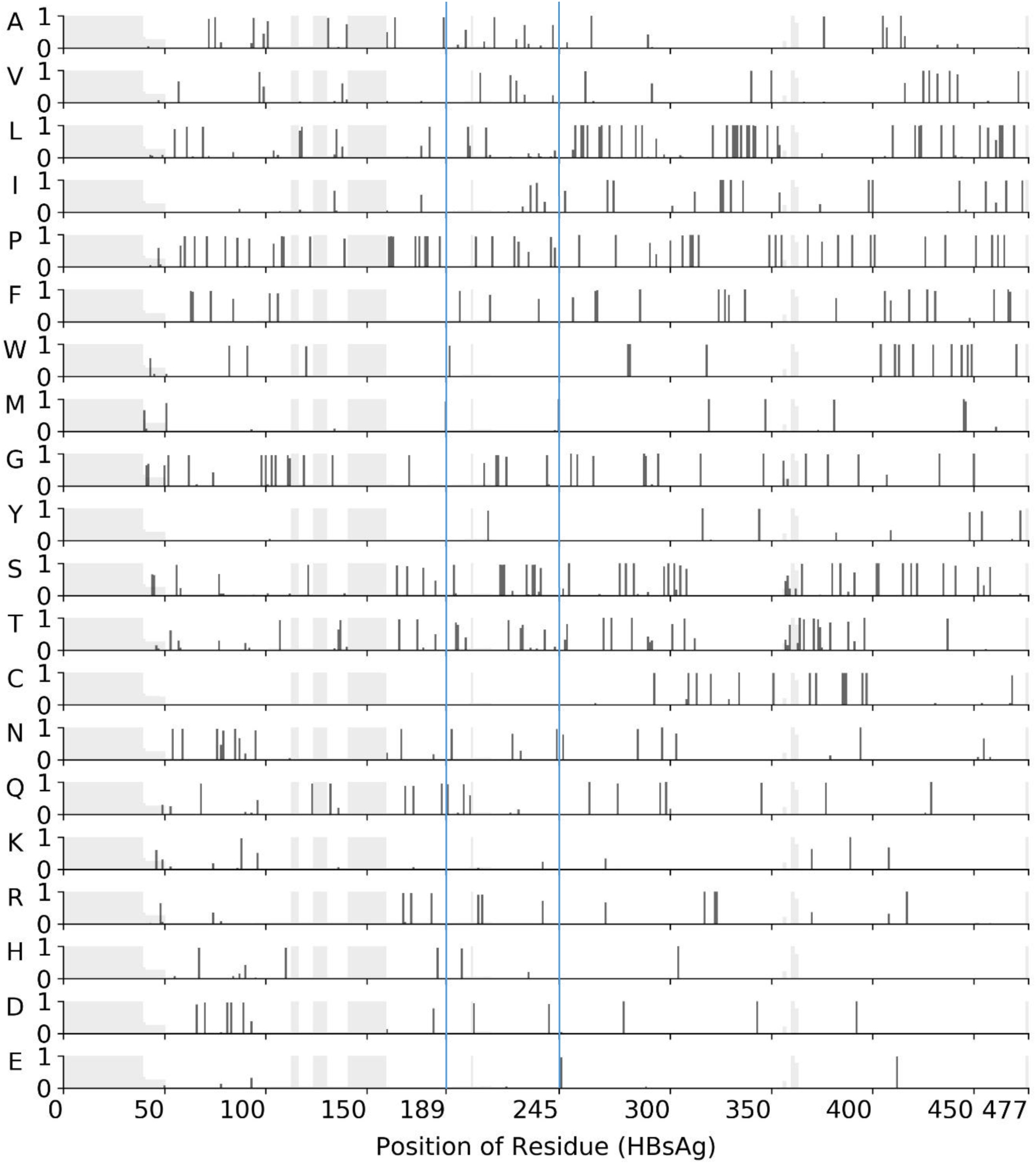
The frequencies of 20 amino acids in different sites of ORF S (the HBsAg). The position (1-188) is the preS1 domain. Position (189-244) is the preS2 domain, position (245-477) is the S domain.

**Fig. S7.**
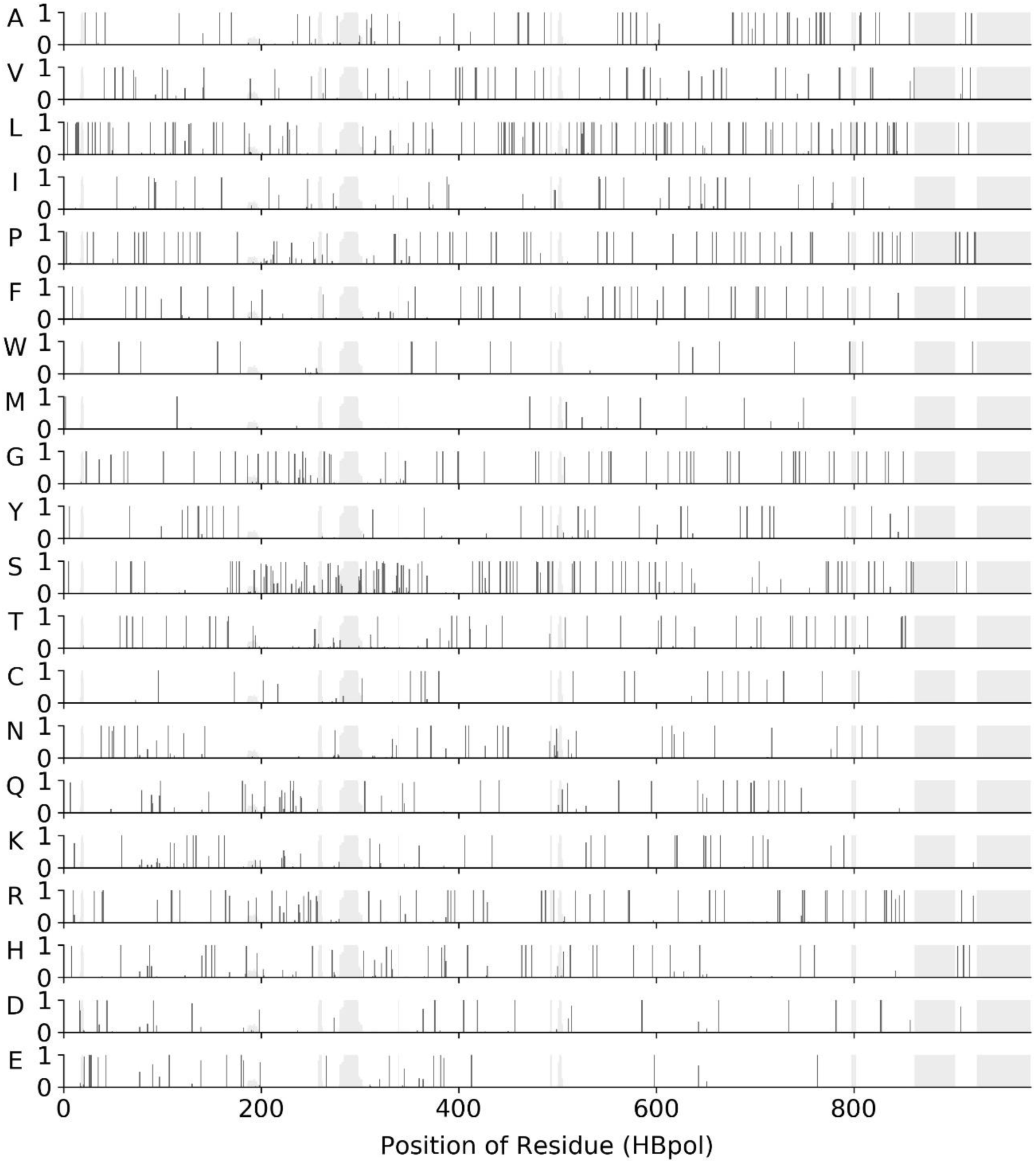
The frequencies of 20 amino acids in different sites of ORF P.

**Fig. S8.**
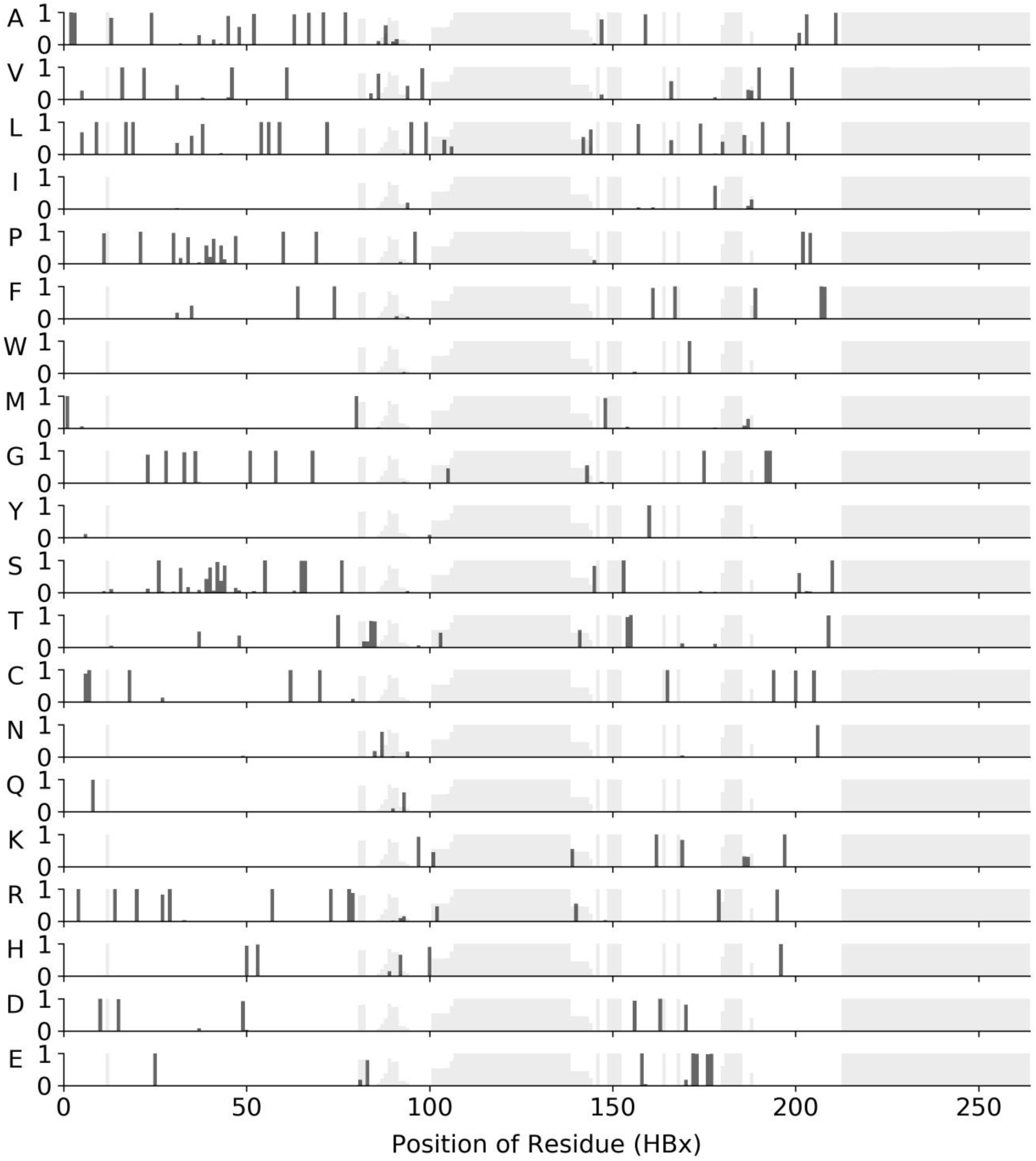
The frequencies of 20 amino acids in different sites of ORF X.

